# Development and optimization of a high-throughput 3D rat Purkinje neuron culture

**DOI:** 10.1101/2020.05.20.105858

**Authors:** Ida M. Uggerud, Torbjorn Krakenes, Hirokazu Hirai, Christian A. Vedeler, Manja Schubert

## Abstract

Improved understanding of the mechanisms involved in neurodegenerative disease has been hampered by the lack of robust cellular models that faithfully replicate *in vivo* features. Here, we present a refined protocol for generating age-dependent, well-developed and synaptically active rat Purkinje neurons, responsive to paracrine factors and supporting a 3D cell network. Our model provides high experimental flexibility, high-throughput screening capabilities and reliability to elucidate Purkinje neuron function, communication and neurodegenerative mechanisms.

## Article

Unravelling the mechanisms of neurodegeneration depends on the availability of robust models that provide insight both at the single cell level and network levels, and that offer high experimental flexibility. Dissociated neuronal cultures can be useful, but their quality and survival dependents on several factors including animal species, age of tissue that is dispersed to give single cells, the surface onto which the single cells are seeded and cultured, and co-factors that drive neuronal growth and development. To date, the majority of successful Purkinje neuron cultures (PNC) models have used embryonic mouse cerebellum, few have successfully used rat embryonic or postnatal cerebellum. Although the success rate of transgenic alterations and *in vivo* modelling is lower in rats ^1^, the rat is physiologically, genetically and morphologically closer to humans than the mouse ^2^, and outbred or transgenic rat models mimic human neurodegenerative disease mechanisms and progressions more closely ^3–5^ than mouse models do ^6,7^.

Since neurodegeneration generally occurs in the adult or aged human brain, a dissociated culture system derived from mature rather than embryonic tissue is desirable. However, previous attempts to culture functional dissociated neurons from late postnatal and adult tissue have been largely unsuccessful. Therefore, our goal was to develop a culture protocol that provided well-developed, mature, functional and synaptically active rat Purkinje neurons (PNs), interdependent of the derived tissue age, that gaves maximal experimental flexibility and the potential for high-throughput screening. We discovered three factors that were essential for success: having a three-dimensional (3D) growth structure, pH stability and co-factor supplementation.

The first attempt growing PNs directly on glass cover-slips coated with poly-D-lysine (PDL) and the extracellular matrix protein laminin failed: the yield of PNs per cover-slip declined to zero from E18 to P10 at 21 days *in vitro* (*DIV*) (Figure 1a, non 3D-SCL). We reasoned that the extracellular matrix used lacked important features including other cell types that provide the *in vivo* 3D cell network structure and thereby cell-cell communicate including paracrine factor secretion. Therefore, in the second attempt we developed a three-dimensional support cell layer (3D-SCL) approach by plating two cerebellar cell layers derived of either E18, P0 or P10 tissue onto PDL coated cover-slips. We introduced a time-delay by plating the second cell layer 7 to 48 days later than the first. We found that the tissue age of cells used to grow the 3D-SCL (E18 to P10) had no impact on the PN yield of the second layer, however, there was a strong correlation between the *in vivo* age of the support cell layer and the tissue age used to grow the second cell layer, the enriched PN layer. The highest survival rate of E18 derived-PNs was observed when plated onto the 3D-SCL at *DIV14*, for P0 derived-PNs at *DIV21* and for P10 derived-PNs at *DIV28* (Figure 1a). These findings indicate that the older the starting tissue, the more mature the 3D-SCL has to be to achieve a high survival rate of PNs for a minimum of 21 to 28 *DIV*.

**Figure 1.**
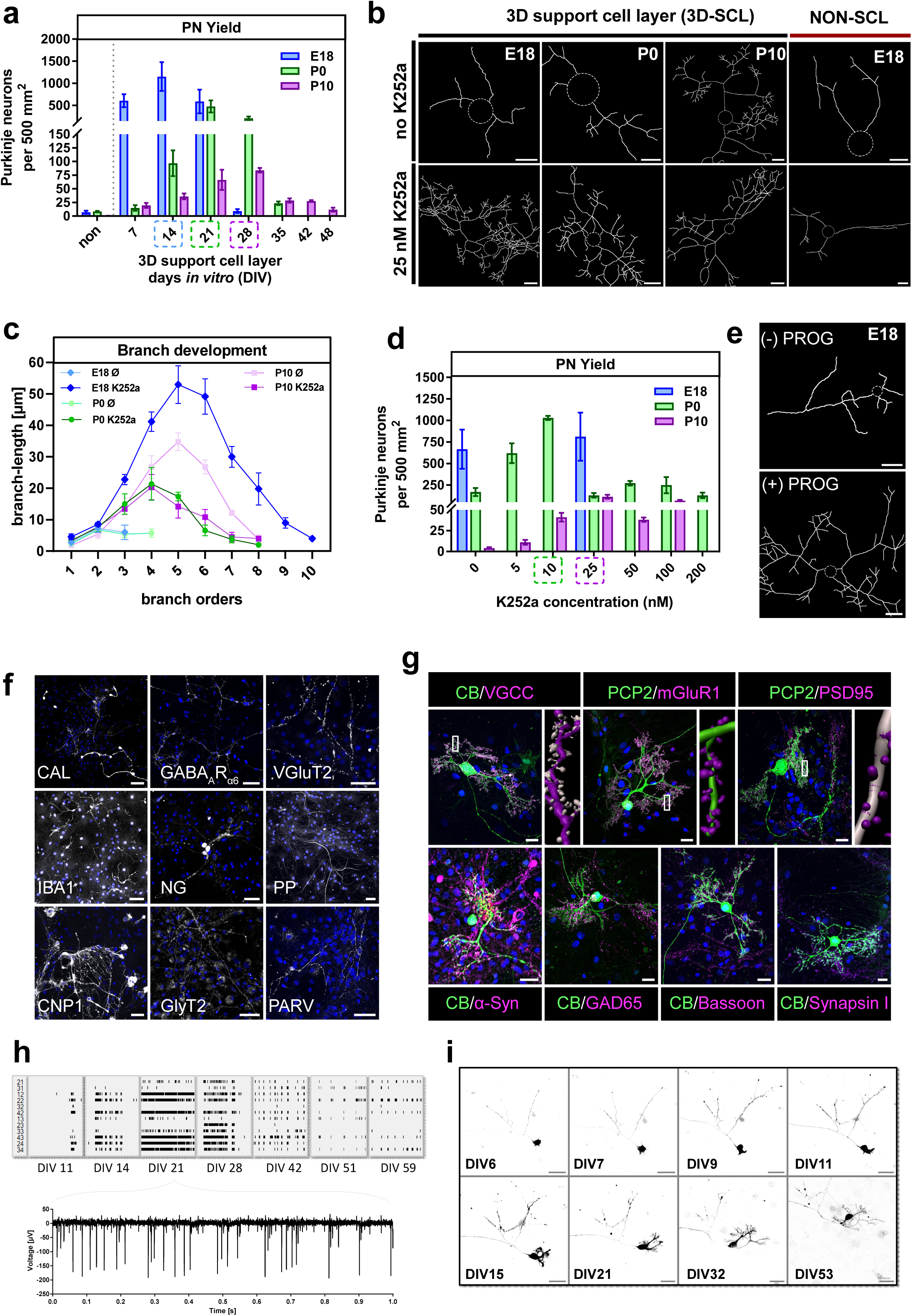
Evaluation of age-dependent rat Purkinje neuron culture. (**a**) Interdependent relationship of Purkinje neuron yield and *in vitro* age of the 3D support cell lager (3D-SCL: DIV 7 to 48) for E18, P0 and P10 derived-Purkinje neurons. (**b**) Representative Purkinje neuron skeletons dependent on derived neuron age, 3D-SCL and protein kinase C (PKC) antagonist K252a. Scale bar, 20 µm; (**c**) Analysis of dendritic branch structure towards length and branch orders for Purkinje neurons derived from E18, P0 and P10 tissue without and with 25 µM K252a to modulate PKC activity. (**d**) Interdependent relationship of Purkinje neuron yield and concentration-dependent PKC activity modulation for E18, P0 and P10 derived-Purkinje neurons. (**e**) Representative skeleton of an E18 derived-Purkinje neurons visualizing the effect of 40 µM progesterone on dendritic branching. Scale bar, 20 µm; (**f**) Immunohistochemical representation of the major cell types (white) forming the 3D-SCL: unipolar brush cells (CAL-calretinin), granule cells (GABAARα6), Golgi cells (NG-neurogranin, GlyT2), Lugaro cells (GlyT2), stellate and basket cells (PAV-parvalbumin), fibres such as mossy and climbing (VGluT2, PP-peripherin), oligodendrocytes (CNP1) as well as microglia (IBA1). Nuclei staining DAPI (blue). Scale bar, 50 µm; (**g**) Immunohistochemical representation of mature Purkinje neurons (green; CB-calbindin, PCP2 - Purkinje cell specific protein 2) positive for post- and presynaptic biomarkers (magenta). Postsynaptic: VGCC, mGluR1, and PSD95 including 3D IMARIS cartoon reconstruction of the protein positive synapses on one chosen Purkinje neuron dendrite; Pre-synaptic: α-synuclein (α-syn) – marker of glutamatergic synaptic terminals from granule cells (parallel fibres) and unipolar brush cells (type I/II); GAD65-marker of axon terminals from stellate and basket cells; bassoon – marker of the active zone of mossy fibre terminals and parallel fibre terminals between Golgi cells and granule cells, and between basket cells and Purkinje neurons; and synapsin I – synaptic vesicle phosphoprotein of mature CNS synapses; Nuclei staining DAPI (blue). Scale bar, 20 µm; (**h**) MEA recorded spike patterns (10s) with a cut-out (1s) at day 21 *in vitro* following Purkinje neuron maturity. (**i**) Live-cell imaging of E18 derived-Purkinje neuron expressing lentiviral-induced GFP from day of seeding (DIV0) up to 2 months (DIV53). The Purkinje neuron development to maturity was very similar to *in vivo*, as the fusion phase (E17 - P5 ≈ *DIV0 – DIV7*), the phase of stellate cells with disoriented dendrites (P5 - P7 ≈ *DIV7 – DIV9*), as well as the phase of orientation and flattering of the dendritic tree (P7 - P21 ≈ *DIV9 – DIV23*) were observed. Scale bar, 50 µm

However, the use of a “double” cell layer was associated with higher metabolic demand than single layer cultures and led to non-physiological pH fluctuations resulted in cell death when half of the culture media was replaced ones a week. Replacing the culture media more frequently, either every 3.5-days (6 well) or every 2-days (12 and 24 well) prevented pathological pH fluctuations and gave a healthy well-developed neuronal network. Despite this, immunofluorescent staining showed that the PNs had a poorly developed dendritic morphology compared to those *in vivo*, with fewer and shorter branches in E18 and P0 derived-PNs (Figure 1b-c, 1b upper panel).

Neuronal dendrites are generated during development by a series of processes involving a first step of extension and retraction of dendritic branches, and subsequently stabilisation of existing dendrites through building of synaptic connections and neuronal calcium homeostasis ^8^. Calcium-dependent protein kinase C (PKC) subtypes, activated by synaptic inputs from the parallel fibres (granule cells) through metabotropic glutamate receptors (mGluR1/4), trigger functional changes as well as long-term anatomical maturation of the PN dendritic tree during cerebellar development ^9^. Altering the activity of calcium-dependent PKC subtypes using PKC antagonist K252a improved dendritic branching for E18 and P0 derived-PNs similar to *in vivo*, but had no effect on the branching characteristics of P10 derived-PNs (Figure 1b-c, 1b lower panel). Interestingly, K252a-induced PKC inhibition significantly improved the low survival rate observed for P0 and particularly for P10 derived-PNs in a concentration dependent manner (Figure 1d). The survival rate in P0 derived-PNs was improved by a factor of 6 by blocking 20 % of PKC activity (10 nM K252a), whereas in P10 derived-PNs, blocking PKC activity to 50 % (25 nM K252a) increased the survival rate by a factor of 28. Inhibiting PKC activity had no effect on the survival rate of E18 derived-PNs (Figure 1d).

PN survival and dendritic tree development are also highly dependent on paracrine factors such as progesterone, insulin and insulin-like growth factor 1 (IGF1) secreted by other cells or self-produced by PNs in an age-dependent manner ^10–12^. We supplemented our culture with 40 µM progesterone and found this led to increased branched dendritic trees in E18 derived-PNs, but it had no impact on the branch structure of P0 and P10 derived-PNs (Figure 1e). Even though PN dendritic development was insufficient when either K252a inhibition or progesterone were not supplied, supplementation with insulin and IGF1 were sufficient to maintain the long-term growth of the other cerebellar cell types: granule, Golgi, Lugaro, unipolar brush, stellate and basket cells (Figure 1f).

To demonstrate that our PNs expressed functional synapses, we used immunocytochemistry to identify pre- and postsynaptic biomarkers of functional synapses including voltage-gated calcium channels (VGCC), metabotropic glutamate receptor 1 (mGluR1), post-synaptic density protein 95 (PSD95), glutamate-decarboxylase 65 (GAD65), glycine transporter 2 (GlyT2), α-synuclein and bassoon. All these markers were present indicating a level of maturity of both the PNs and the surrounding network (Figure 1g).

Next, we tested the functional activity of these PNs. *In vivo*, PNs fire spontaneous action potentials at frequencies of about 40-50 Hz with a complex trimodal pattern of tonic firing, bursting, and silent modes that depend on anatomically and functionally maturity ^13,14^. E18 derived-PNs cultured in a 24 well multielectrode array first revealed spontaneous bioelectrical activity on *in vitro* day 11. The spike rate increased constantly from 0.15 ± 0.03 Hz (*DIV11*) to 2.56 ± 0.59 Hz (*DIV21*). After *DIV*28, the spike activity become erratic with long periods of silence, but overall, a frequency of 2.79 ± 0.55 Hz was maintained until *DIV*63 (Figure 1h). We observed uniform, highly non-uniform spike intervals and trains with silent periods between bursts and spike frequencies of up-to 140 Hz within the burst. Exchanging the PNC media at *DIV28* to one previously used in organotypic brain slice culture^15^, prevented the erratic spike activity and stabilized the spike frequency at 6.35 ± 1.85 Hz for up-to 63 *DIV*.

In addition to immunocytochemical and high-throughput electrophysiological studies, this 3D PN model system will provides the potential for cell-type-specific genetic engineering. For example, by using lentiviral particles to express PN-specific green fluorescence protein (GFP) via implementation of the L7 promoter ^16,17^. To test this, we applied L7-GFP inducing viral particles to dissociated PNs on the day of seeding. Within 3 days, we found PNs expressing GFP and hardly any off-targets (<0.02%). At *DIV14*, 61.5 % of the PN population were GFP positiveand these cells did not differ in dendritic structure and stably expressed GFP for up-to 169 *DIV* (Figure 1i). Using our culture system, we also found a sufficiently high transfection rate of PNs when lentiviral particles were added to the culture at *DIV14* and *DIV28*, however the rate of transfection and speed of expression fell progressively the later the genetic manipulation was implemented. The GFP positive PNs in the culture revealed a very similar development to *in vivo*, as we were able to observed the fusion phase (E17-P5), the phase of stellate cells with disoriented dendrites (P5-P7), as well as the phase of orientation and flattering of the dendritic tree (P7-P21) ^18,19^ (Figure 1i).

We present a 3D, rat PNC model for growing Purkinje neurons that is independent of derived tissue age, and which provides a complex and robust system that allows maximal experimental flexibility. The combined use of 3D network structures (3D-SCL) with optimized concentrations and time-dependent addition of hormones, paracrine factors and activity regulators (progesterone, insulin, IGF-1, K252a), created ideal conditions to grow a balanced cerebellar network in miniature (Figure 2). As a proof-of-principle, we demonstrated the usefulness of this culture model as a high-throughput screening tool to investigate disease mechanisms including drug/compound testing. The long-term stability and neuronal complexity of our culture will facilitate the study of cell- and network-dependent cerebellar degeneration related to paraneoplastic cerebellar degeneration and ataxia.

**Figure 2.**
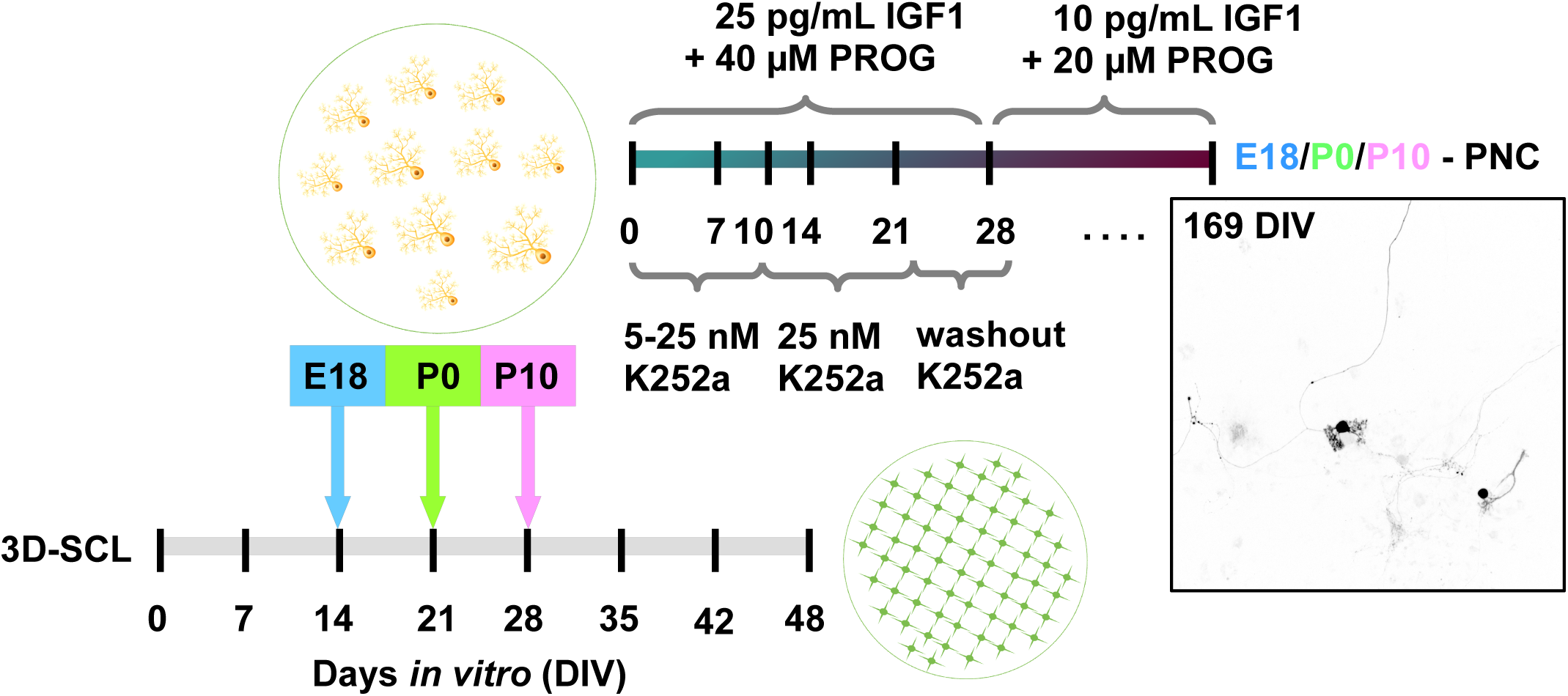
Optimized 3D rat Purkinje neuron culture protocol. Each tested culture desired different conditions of support and activity interdependent of the starting tissue age. Whereas the supplementation of insulin-like growth factor 1 (IGF1) and progesterone (PROG) induced a stable environment to obtain high survival rates of Purkinje neurons, PKC activity modulation mainly shaped the dendritic tree development, with the exception of P10 tissue derived neurons where the survival was highly dependent on the inhibition of PKC but not their dendritic tree development. The optimized protocol for all tested tissues relies on the time point of placing the second cell layer, the Purkinje neuron enriched layer, and media that is supplemented with IGF1, progesterone and K252a, where K252a starting concentration is altered dependent on the used tissue to start the culture as follow; DIV1-10: E18 - 5 nM, P0 - 10 nM, P10 - 25 nM; DIV10-22: the K252a concentration is raised to 25 nM for E18 and P0 until the dendritic tree is well-developed and mature; DIV22-28: washout phase, K252a supplementation is stopped (DIV22-24: 12.5 nM, DIV24-26: 6.75 nM, DIV26-28: 3.35 nM). At DIV 28 the IGF1 and progesterone concentration is reduced by factor, 2.5 and 2, respectively, to proceed to long-term culture conditions. The developed protocol allows to grow a stable Purkinje neuron 3D culture for up to 6 months (DIV163) in a 6 to 24 well format.

## Material and Methods

### Neuronal culture preparation

All procedures were performed according to the National Institutes of Health Guidelines for the Care and Use of Laboratory Animals Norway (FOTS 20135149/20157494/20170001). Wistar Hannover GLAST rat pups (n = 328), embryonic day 18 (E18) to postnatal day 10 (P10), were used for neuronal culture preparation.

Briefly, following anaesthesia and decapitation, the brains were rapidly transferred into preparation solution: ice-cold EBSS solution (Gibco, #24010043) containing 0.5% glucose (Sigma, #G8769) and 10 mM HEPES (Gibco, #15630056). Under a dissection microscope, carefully remove the meninges, cut off the medulla oblongata and separate the cerebellum from the pons and the midbrain. Depending on the culture, Purkinje neuron or structural layer, transfer either only the cerebellum or the cerebellum including pones to a 15 mL tube containing 20 U/mL papain (Worthington, #LK003178) solved in preparation solution and warmed up to 36 °C. Place the tube into the incubator for 15 minutes at 36°C with occasionally swirling to digest the tissue. Remove the papain solution carefully with a fire polished Pasteur pipette and stop the digestion by adding pre-warmed stop media (36°C): advanced DMEM/F12 solution (Gibco, #12634010) containing 0.5% glucose (Sigma, #G8769) and 10% foetal bovine serum (FBS, Gibco, #10500064). After 5 minutes of deactivation, remove the stop media and add 250 µL growth media containing 10% FBS per cerebellum and pipette the tissue/media suspension with a fire polished Pasteur pipette 100X until cells are separated.

### 3D Support Cell Layer (3D-SCL)

375000 cells/mL from cerebellum including pones were seeded on pre-coated coverslides from Neuvitro (#GG-12-1.5-PDL, 24 well, 500 µL/well; #GG-18-1.5-PDL, 12 well, 1 mL/well; #GG-25-1.5-laminin, 6 well, 2 mL/well). Culture were maintained in 6-,12- or 24-well plates in growth media consisting of 45% advanced DMEM/F12 solution (Gibco, # 126340010), 45% NBM solution (Miltenyibiotec, #130-093-570), 1.5% B-27 serum-free supplement (Gibco, #17504044), 1.5% NB-21 serum-free supplement (Miltenyibiotec, #130-093-566), 1% NaPyruvate (Invitrogen, #11360088), 1% heat-inactivated FBS (Invitrogen, #10500064), 2% GLUTAMAX (Gibco, #35050038), 5 mg/mL D-glucose and 10 mM HEPES (Invitrogen, #15630056) at 36°C. Half of the culture medium was replaced every 7 days.

### Purkinje neuron layer

E18 and P0 derived Purkinje neuron culture: 500000 cells/mL from cerebellum without pones were seeded on the 3D support cell layer of different *in vitro* ages. P10 derived Purkinje neuron culture: 750000 cells/mL from the vermis of the cerebellum were seeded on the 3D support layer of different *in vitro* ages. The growth media was supplemented with insulin (Invitrogen, #12585014; 1:250, stock 4 mg/mL), progesterone (Sigma, #P8783, 1:2000, stock 80 mM), insulin-like growth factor 1 (IGF1; Promokine, #E-60840, 1:40000, stock 1 µg/µL) and Protein kinase C inhibitor K252a (Alomone, # K-150; IC_50_ 25 nM). In long-term cultures that were maintained for more than 28 days *in vitro* the IGF1 and progesterone concentration were reduced to 10 ng/mL and 20 µM, respectively. K252a was supplemented for 21 days before the washout process started, its optimal concentration was experimental evaluated for each tested culture type. Half of the culture medium was replaced every 3.5 (6 well) and 2 (12/24 well) days, respectively. All experiments testing the Purkinje neuron yield dependent on derived tissue age, *in vitro* age of the 3D-SCL and K252a concentration were performed randomly, containing 3 to 6 probes per experimental setting and 5 independently repeats for each group and condition.

### Lentiviral gene editing

L7 promoter (full length 1005 bp) were custom cloned by SBI System Bioscience into construct pCDH-L7-MCS-copGFP (#CS970S-1) and viral particle with a yield of 2.24 × 10^9^ ifus/mL were produced. Freshly prepared Purkinje neurons of E18 or P0 cerebellum suspended in growth media containing no serum were incubated for 10 minutes at 37 °C with 1.22 × 10^6^ viral particle/mL before seeded onto the supplement structure layer containing cover-slip or live cell imaging µ-dish (#80136, 35 mm, Ibidi). Media was changed after 3 days and transfection efficiency evaluated by live cell imaging microscopy 24h post transfection, daily up to 21 days and weekly up to 169 days in culture, respectively. Additional, lentiviral transfection of Purkinje neurons in culture were performed 1 day after feeding at DIV15 and DIV29 by applying 2.5 × 10^6^ viral particle/mL to evaluate the efficiency and effects of age-dependent genetic manipulations. The neuronal development of the GFP expressing Purkinje neurons was followed by obtaining 10 independent 3×3 tile scan using the Zyla camera configuration (2048×2048) with the CFI Plan Apochromat Lambda dry objective 10×0.45 (pixel size 603 nm) or 20×0.75 (pixel size 301 nm) at the Andor Dragonfly microscope system (Oxford Instruments company). The experiments of DIV0, DIV15 and DIV29 were repeated three times.

### Immunohistochemical cell type characterisation

To evaluate Purkinje neuron yield and the distribution ratio of other cell types of the cerebellum, including their synaptic interactions, the culture was washed with pre-warmed 0.1 M PBS (1xPBS; Gibco, #70013016) and fixed with 1.5-4% paraformaldehyde (PFA, pH 6-7.2; ThermoScientific, #28908) containing 0.5% sucrose for 15 minutes at 36°C. Tris-based or citric acid-based heat induced antigen retrieval (pH 9 and pH 6; 45 min, 85 °C) ^20^ were perform when necessary (see Table 1). Culture were quenched with 1xPBS containing 50 mM NH_4_Cl (PBS_N_), permeabilised with 0.2% Triton X-100 (Sigma, #T9284) in PBS_N_ (5 min, 36°C), rinsed with PBS_N_ containing 0.5% cold water fish gelatine (Sigma, #G7041)(PBS_NG_, 3×15 min), and incubated with primary antibody over-night at 4°C in PBS_NG_ containing 10% Sea Block (SB; ThermoScientific, #37527), 0.05% Triton X-100 and 100 μM glycine (Sigma, #G7126) to visualise the different cerebellar cell types, including Purkinje neurons and their synaptic interactions (Table 1). The cover-slips were rinsed with PBS_NG_ (3×20 min) and incubated with highly cross-absorbed donkey secondary antibodies conjugated to CF™488/594/647-Dye (1:400; Biotium, #20014, #20115, #20046, #20015, #20152, #20047, #20074, #20075, #20169, #20170) for 2 hours at 22°C in PBS_NG_ containing 2.5% SB. To remove unbound secondary antibody cover-slips were rinsed with PBS_N_ (3×20 min), and briefly tipped into MilliQ water before mounted in hardening Prolong™ Glass Antifade Reagent (Invitrogen, #P36981) onto cover-slides. After 2 days of hardening at 18-21°C in the dark, cover-slides were stored at 4°C until imaging.

**Table 1.**
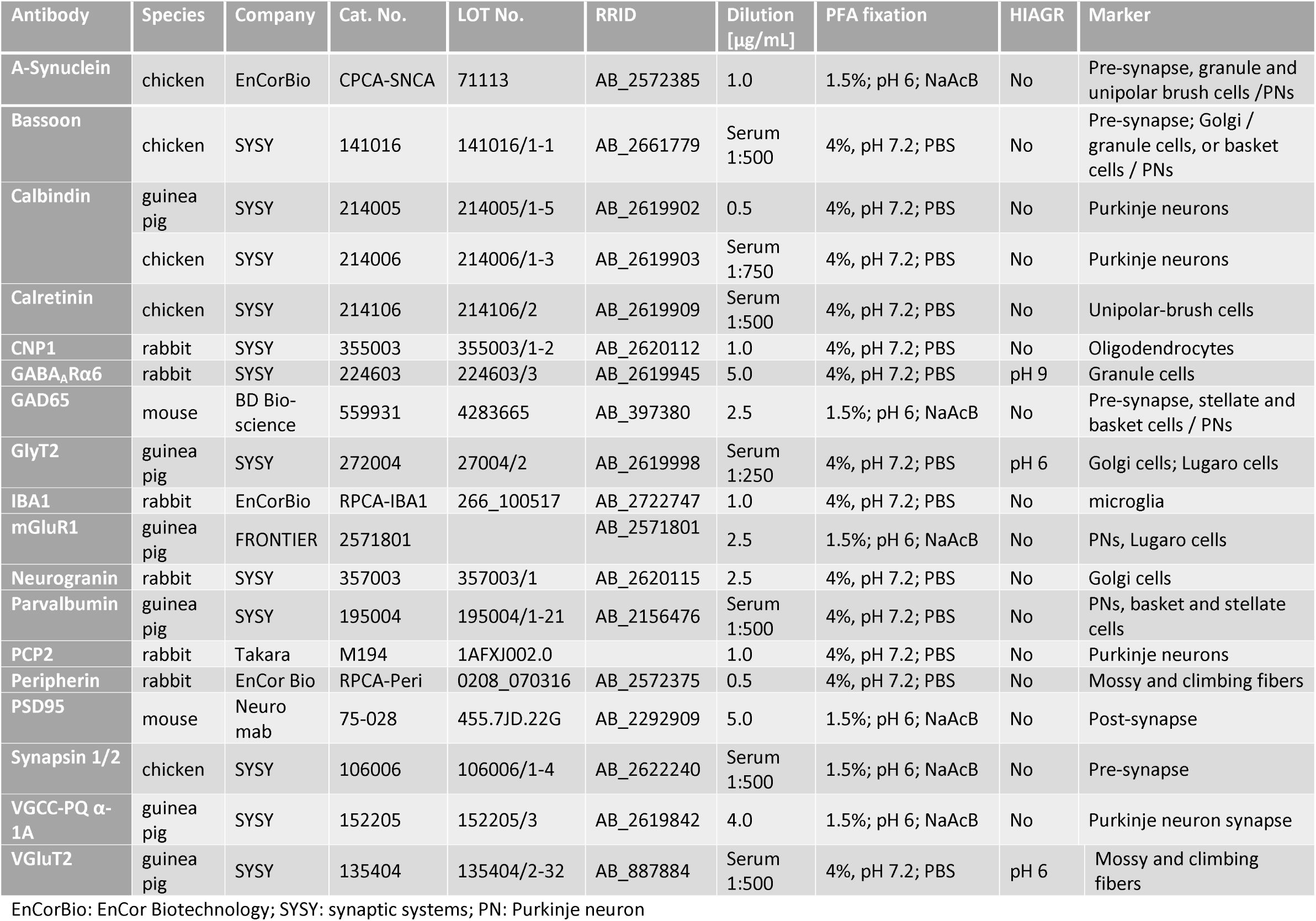
Primary antibodies. The signal to noise ratio for the antibodies were evaluated for the following conditions: 4% PFA at pH 7.2 diluted in 100 mM PBS; 1.5% PFA at pH 6 diluted in 100mM natrium acetate buffer (NaAcB)); without heat-induced antigen retrieval (HIAGR); and with HIAGR either TRIS-based (pH 9) or citric acid-based (pH 6). The best conditions for each used antibody are described below.

### Purkinje neuron count and imaging

Purkinje neurons were counted manually and blind by screening the cover-slips using a Leitz Diaplan Fluorescence microscope equipped with CoolLED pE-300white. For dendritic tree branch analysis and determination of maturity and synaptic interaction, 10 Purkinje neuron Z-stack images per cover-slide were collected in 5 independent and randomized experiments at 0.5-1 μm intervals with the Zyla camera configuration (2048×2048) at the Andor Dragonfly microscope system using either a CFI Plan Apochromat Lambda S LWD 40×1.14 water objective (pixel size 151 nm), 60×1.20 oil objective (pixel size 103 nm) or CFI SR HP Apo TIRF 100×1.49 oil objective (pixel size 60 nm) to detect DAPI and CF™488/594/647 dye emission and superimposed with Fusion software (Oxford Instruments). 3D surface visualization of synapses was performed using Oxford Instruments analysis software IMARIS 9.3.1 and the filament tracer tool ^21^.

### Dendritic tree branch analysis

The Purkinje neuron dendritic tree development was evaluated by analysing group dependent 10 Purkinje neurons per experiment in 10 independent experiments towards the order and length of the dendritic arbours by using an open-source ImageJ and Fiji plugin Simple_Neurit_Tracer (Neuroanatomy) ^22^.

### Micro-electrode array (MEA) recordings

Primary cultures of E18 derived-PNs at a concentration of 500000 cells/mL were plated onto PDL precoated 24 well format plate of the Multiwell-MEA-system (Multi Channel System-MCS, Reutlingen, Germany). Each well contains 12 PEDOT coated gold micro-electrodes (30 µm diameter, 300 µm space, 3 × 4 geometrical layout) on glass base to facilitate visual checking (#890850, 24W300/30G-288). The amplifier (data resolution: 24 bit; bandwidth: 0.1 Hz to 10 kHz, modifiable via software; default 1 Hz to 3.5 kHz; sampling frequency per channel: 50 kHz or lower, software controlled; input voltage range: ± 2500 mV), stimulator (current stimulation: max. ± 1 mA; voltage stimulation: max. ± 10 V; stimulation pattern: pulse or burst stimulation sites freely selectable) and heating element (regulation: ± 0.1 °C) is integrated in the Multiwell-MEA-headstage which is driven by the MCS-Interface Board 3.0 Multiboot. The Multiwell recording platform is covered by a mini incubator to provide 5% CO_2_ and balanced air. Electrophysiological signals were acquired at a sampling rate of 20kHz through the commercial software Multiwell-Screen. Plates were tested every second day for spontaneous activity from day 5 *in vitro*. Raw voltage traces were recorded for 120 seconds, saved and analysed using offline MCS-Multiwell-Analyzer to calculate spike rate and burst activity, including network properties. Two experimental settings were tested: number 1 recording of spontaneous spike activity in Purkinje neuron culture media (45% advanced DMEM/F12 solution, 45% NBM solution, 1.5% B-27 serum-free supplement, 1.5% NB-21 serum-free supplement, 1% NaPyruvate, 1% heat-inactivated FBS, 2% GLUTAMAX, 5 mg/mL D-glucose, 10 mM HEPES, 16 µg/mL insulin, 25 ng/mL IGF1, 40 µM progesterone, 5 nM K252a) for 63 days and number 2 recording spontaneous spike activity for the first 28 days in Purkinje neuron culture media but then exchanged to organotypic brain slice culture media ^15^ (30% advanced DMEM/F12 solution, 20% MEM solution (#41090028; Gibco), 25% EBSS solution (#24010043; Gibco), 25% heat-inactivated horse serum (#H1138; Sigma), 2% GLUTAMAX, 5 mg/ml D-glucose and 2% B-27 serum-free supplement) for the remaining 45 days.

### Notes to provide stable high yield Purkinje neuron culture

1. All media should be prepared fresh on the day of use.
2. Prevent repeated thaw-freeze cycles of the supplements
3. 3D-SCL should be fed 24 hours prior plating of the second cell layer, PN layer, to provide stable pH at 6.8 to 7.0 on the day of seeding.

## ACKNOWLEDGMENT

The authors thank Y.Ishizuka and C.E.Bramham for providing the E18 cerebellum tissue, C. Elliott for discussion, and the Molecular Imaging Centre (MIC), where the imaging experiments were performed (Department of Biomedicine and the Faculty of Medicine and Dentistry of University of Bergen). This work was funded by grants from HelseVest Norway and University of Bergen.

## AUTHOR CONTRIBUTIONS

M.S. devised the conceptual framework. I.M.U., T.K and M.S planned and performed the experiments and analysed the obtained data sets. H.H. provided the lentiviral approach. The paper was written by M.S, H.H. and C.A.V. with editing contributions from all the authors.

